# Rapid screening of target-specific isotope labeling protocols for *in situ* NMR of bacterial membrane proteins

**DOI:** 10.64898/2026.11.14.766694

**Authors:** Nicholas A. Wood, Tata Gopinath, Francesca M. Marassi

## Abstract

We describe an approach for the efficient development of targeted isotope labeling protocols leading to nuclear magnetic resonance (NMR) structural studies of bacterial membrane proteins in intact cells and native outer membrane (OM) vesicles (OMVs). The assay relies on the bioorthogonal non-canonical amino acid tagging (BONCAT) approach for metabolic incorporation of an azide-bearing residue, followed by copper-catalyzed azide-alkyne cycloaddition (CuAAC) to an alkyne-functionalized fluorescent dye. Fluorescence detection in SDS-PAGE provides a rapid way to test samples for off-target background signal from endogenous proteins, bypassing the need for intensive NMR screening experiments. Using this approach, we develop an optimized protocol that enables high-resolution NMR spectra to be obtained *in situ* with negligible background signal. The BONCAT approach also enables target-specific fluorescence or nano-gold labeling of proteins at the OM surface, further extending the utility of this protocol for correlative structure-activity studies of membrane proteins *in situ*.

## INTRODUCTION

Targeted labeling strategies are critical for structural studies of biomolecules in native cellular samples. In the case of *in situ* nuclear magnetic resonance (NMR), targeted isotope labeling – typically ^15^N, ^13^C and ^2^H – is essential for suppressing off-target background NMR signals from endogenous cellular proteins ^1^. For bacteria, the strategy relies on T7/DE3-mediated expression of the plasmid-encoded target protein, coupled with suppression of chromosomal gene transcription and endogenous protein production by the antibiotic rifampicin, a potent inhibitor of RNA polymerase ^2^. Cells are first grown in unlabeled minimal media and the isotopes are added only after rifampin treatment and induction of recombinant gene expression ^3^, but even with this protocol, off-target labeling can linger and thus contribute background NMR signal. For solid-state NMR of bacterial outer membrane (OM) proteins, *E. coli* strains deleted for the major OM proteins OmpA,C,F and LamB can improve target specificity ^4^ but this approach greatly weakens the cell envelope, partly diminishing the impact of working *in situ*.

Here, we describe a cell culture protocol that improves targeted isotope labeling specificity for *in situ* NMR of bacterial OM proteins in native cell membranes, as well as an assay, based on fluorescence detection of bioorthogonal non-canonical amino acid labeling, that facilitates labeling protocol optimization.

## RESULTS

Recently, we showed ^5^ that native OM vesicles (OMVs) can be used directly for NMR structural analysis of the *Salmonella* OM protein PagC, a key promoter of OM vesiculation, under conditions of low pH, Ca^2+^ and Mg^2+^ ion starvation, or in the presence of cationic antimicrobial peptides ^6^. OMVs are biogenically secreted nanoscale spherical structures that originate from the cellular OM and preserve its native highly asymmetric architecture. Working with *Salmonella* PagC, we sought to develop a metabolic labeling protocol that optimizes PagC target specificity for *in situ* NMR.

Considering that the average half-life of *E. coli* transcripts is several minutes ^7^, we reasoned that carryover mRNA remaining after rifampin treatment can allow endogenous protein production to continue for some time after transfer to isotope enriched media, and is the most likely source of off-target labeling. Our protocol optimization strategy focused on minimizing this effect.

Furthermore, to bypass laborious sample screening by NMR detection, and assess labeling efficiency before intensive multidimensional NMR experiments, we developed a screening approach based on bioorthogonal non-canonical amino acid tagging (BONCAT) ^8^ where co-translational metabolic labeling with a non-canonical amino acid bearing an azide reactive group enables the site to be specifically tagged with an alkyne-functionalized fluorescent dye for rapid and sensitive detection by imaging.

Since approximately 4% of the *E. coli* proteome encodes only an N-terminal Met that is removed after synthesis, the non-canonical Met analog L-azidohomoalanine (AHA) provides a unique bioorthogonal reactive handle for copper-catalyzed azide-alkyne cycloaddition (CuAAC). During the metabolic labeling period, all Met residues are replaced with AHA enabling subsequent CuAAC with alkyne-functionalized AlexaFluor-488 (AF488) for fluorescence detection. Prior work ^9^ with nanoscale secondary ion mass spectrometry showed similar levels of either AHA or ^15^NH_3_ incorporation in proteins, demonstrating that BONCAT is a good surrogate for isotope labeling and thus useful for developing NMR labeling protocols.

We tested five metabolic labeling schemes (**Fig. 1A**) with different windows of transcriptional suppression, including pulses of rifampin interleaved by periods of nutrient dissipation or starvation to allow for abundant transcript turnover. Expression of plasmid-encoded PagC (pPagC) and AHA incorporation were analyzed by SDS PAGE of both whole cell lysates and OMVs harvested directly from the cell culture media, and visualized with either Coomassie stain for total protein production (**Fig. 1B, C**) or AF488 fluorescence for AHA incorporation and labeling **(Fig. 1D, E**).

**Figure 1.**
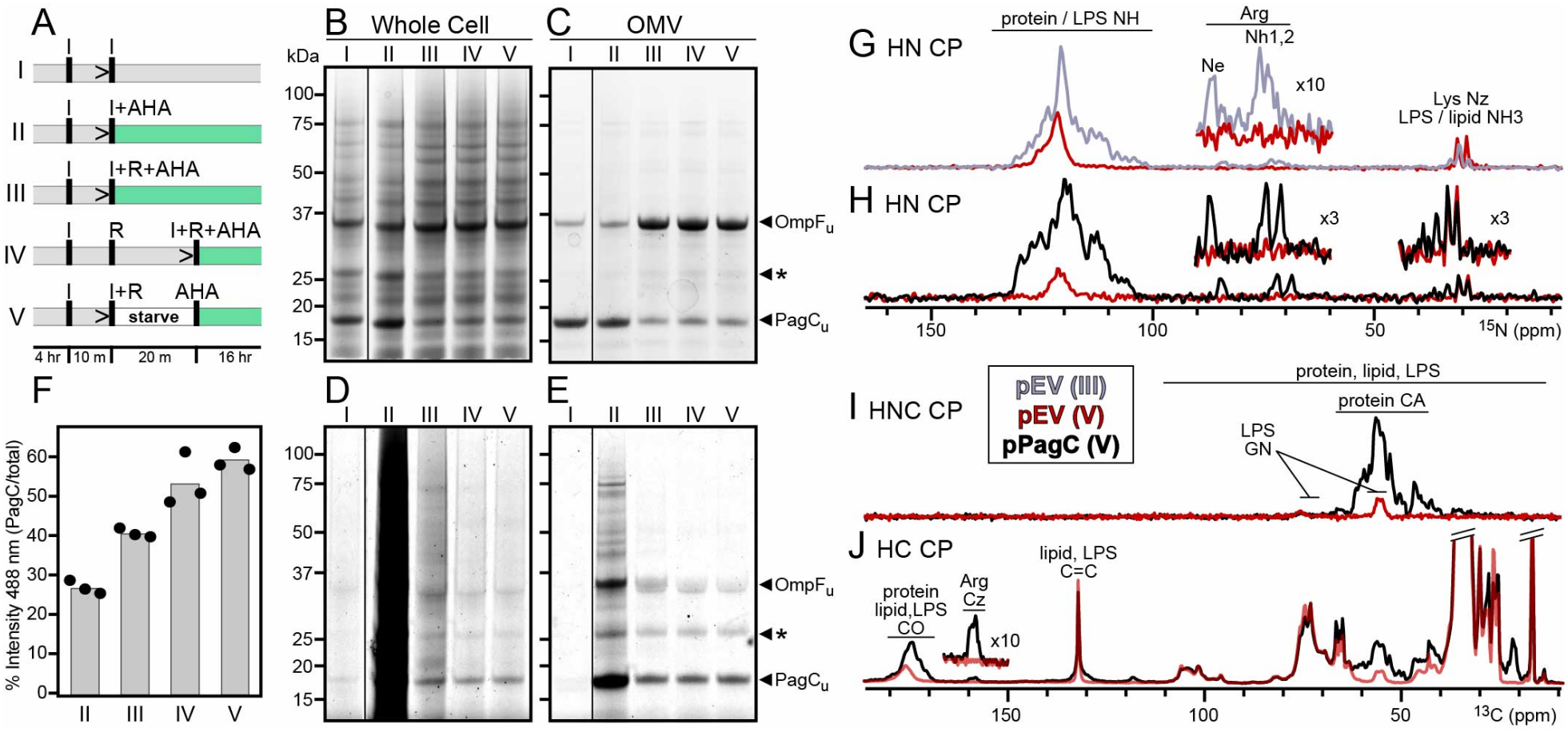
Optimization of targeted labeling. **(A)** Cell culture protocols (I-V) include pulses of IPTG (I), rifampin (R), and the non-canonical amino acid (AHA; green color), media transfers (>), or a period of starvation with no glucose or ammonium salts. **(B-E)** SDS PAGE of total cell lysate (B, D) or OMVs (C, E) after CuAAC with AF488-alkyne. The gels were developed with Coomassie stain (B, C) or imaged with fluorescence at 488 nm (D, E). Bands from OmpF, PagC and a major unidentified protein (*) are marked. **(F)** Relative fluorescence intensity of pPagC-AHA-AF488 OMVs as percent of total intensity. Bar height is the average of three replicate experiments. **(G-J)** Solid-state ^15^N or ^13^C 1D NMR spectra of pEV OMVs produced with protocol III (gray) or protocol V (red), and pPagC OMVs produced with protocol V (black). Spectra were acquired with ^1^H-^15^N (G, H), ^1^H-^15^N-^13^C (I), or ^1^H-^13^C (J) CP, for a total of 10,000(G) or 512 (H-J) transients.

In protocols I and II, cells are grown for 4 hours in label-free minimal media, then treated with two pulses of IPTG separated by 10 minutes in either label-free media (protocol I) or AHA-supplemented media (protocol II), all in the absence of rifampin. Since T7 RNA polymerase is chromosomally-encoded and IPTG-inducible in the *E. coli* BL21(DE3) strain used here, the second pulse of IPTG is essential for all rifampin-treated protocols to ensure that at least one copy is produced before rifampin shuts down endogenous transcription.

As observed previously ^5^, induction with IPTG yields significant levels of pPagC (~18 kDa) in both cells and OMVs (**Fig. 1B, C**) along with the endogenous *E. coli* major porin OmpF (~37 kDa). While pPagC is highly produced in both protocols I and II (**Fig. 1B, C Lanes I, II**), reaction with AF488-alkyne yields abundant fluorescence only when coupled with AHA supplementation (**Fig. 1D, E, Lane II**) and none in its absence (**Fig. 1D, E, Lane I**).

Absent rifampin, all chromosomally-encoded genes are expressed and incorporate AHA. AF488-imaged SDS-PAGE of whole cells cannot distinguish individual protein bands (**Fig. 1D, Lane II**) but OMVs, which only package select proteins reveal approximately equal AF488 intensity for OmpF and pPagC, in addition to a few other proteins (**Fig. 1E, Lane II**). By contrast, the addition of rifampin in protocol III dramatically suppresses endogenous background labeling. This is particularly evident for OMVs where the AF488 fluorescence intensity from OmpF and other non-PagC bands is highly reduced relative to pPagC **(Fig. 1D, E, Lane III**), notwithstanding the high level of OmpF detected by Coomassie stain (**Fig. 1B, C, Lane III**).

Notably, adding a second rifampin pulse 20 min prior to transferring the cells into AHA media (protocol IV) decreases the OmpF AF488 intensity and increases the pPagC target specificity compared to protocol III (**Fig. 1B-E, Lane IV**). Moreover, inserting a 20 min nutrient starvation period, after the rifampin pulse and before transfer to AHA media, further reduces the OmpF AF488 signal and enhances the PagC labeling specificity (**Fig. 1B-E, Lane V**). These results are in line with the expected depletion of transcript RNA during the 20 min period in protocol IV, and more complete degradation of endogenous transcripts during starvation in protocol V.

Quantitative estimates of the pPagC AF488 fluorescence intensity relative to total show that that protocol V provides superior specificity compared to protocol III, which is typically used for *in situ* NMR experiments ^3d^, and to a lesser extent compared to protocol IV (**Fig. 1F**).

To assess targeted isotope labeling for NMR, we used protocols III and V to culture pPagC cells or control cells transformed with empty plasmid vector (pEV) lacking the *pagC* gene, and acquired magic angle spinning (MAS) solid-state NMR spectra of OMVs harvested directly from the culture media, as described ^5^. Since pEV cells do not produce PagC (**Fig. S1**) they provide a useful control for off-target labeling.

The 1D ^15^N spectra, obtained with ^1^H-^15^N cross-polarization (CP) for pEV OMVs from protocol V (**Fig. 1G, red**) reveal a dramatic reduction of ^15^N signal intensity compared to protocol III (**Fig. 1G, gray**). Moreover, while protocol V pPagC OMVs yield abundant ^15^N signal from protein backbone and sidechains (**Fig. 1H, black**), the pEV OMVs have essentially no protein signal, evidenced by the absence of Arg sidechain intensity (**Fig. 1G, H, red**): While the amide NH (~100-140 ppm) and NH_3_ (~35 ppm) regions of the ^15^N spectrum contain signals from protein, lipid and lipopolysaccharide (LPS), the Ne and Nh_1,2_ signals from Arg sidechains (~70-90 ppm) are exclusively from protein, and therefore serve as effective diagnostics of background labeling. The ^15^N spectra thus show that protocol V is superior at suppressing background ^15^N signal from off-target proteins.

The1D ^13^C NMR spectra support this conclusion. The superior background suppression afforded by protocol V is evident in both ^1^H-^15^N-^13^C and ^1^H-^13^C CP spectra (**Fig. 1I, J**) where the pEV OMV spectra show dramatic protein signal reduction compared to pPagC OMVs. Here, the absence of Arg sidechain Cz intensity confirms that the residual background signal is from LPS and lipid and not protein.

In metabolic labeling, AHA is incorporated at all protein Met sites during translation, and the resulting cells and OMVs must be denatured to expose cryptic intracellular or intramembrane AHA and enable CuAAC with AF488-alkyne.

Indeed, reaction with AF488-alkyne in intact whole cells does not produce fluorescence (**Fig. 2A, B**), notwithstanding the abundant expression of pPagC and in line with the intramembrane occluded positions of its two native Met residues (**Fig. 2C**). The Met residues of OmpF and the other major OM proteins are also occluded (**Fig. 2D, Fig. S2**) and thus not expected to be accessible for CuAAC in intact membranes, as indeed observed (**Fig. 2B**).

**Figure 2.**
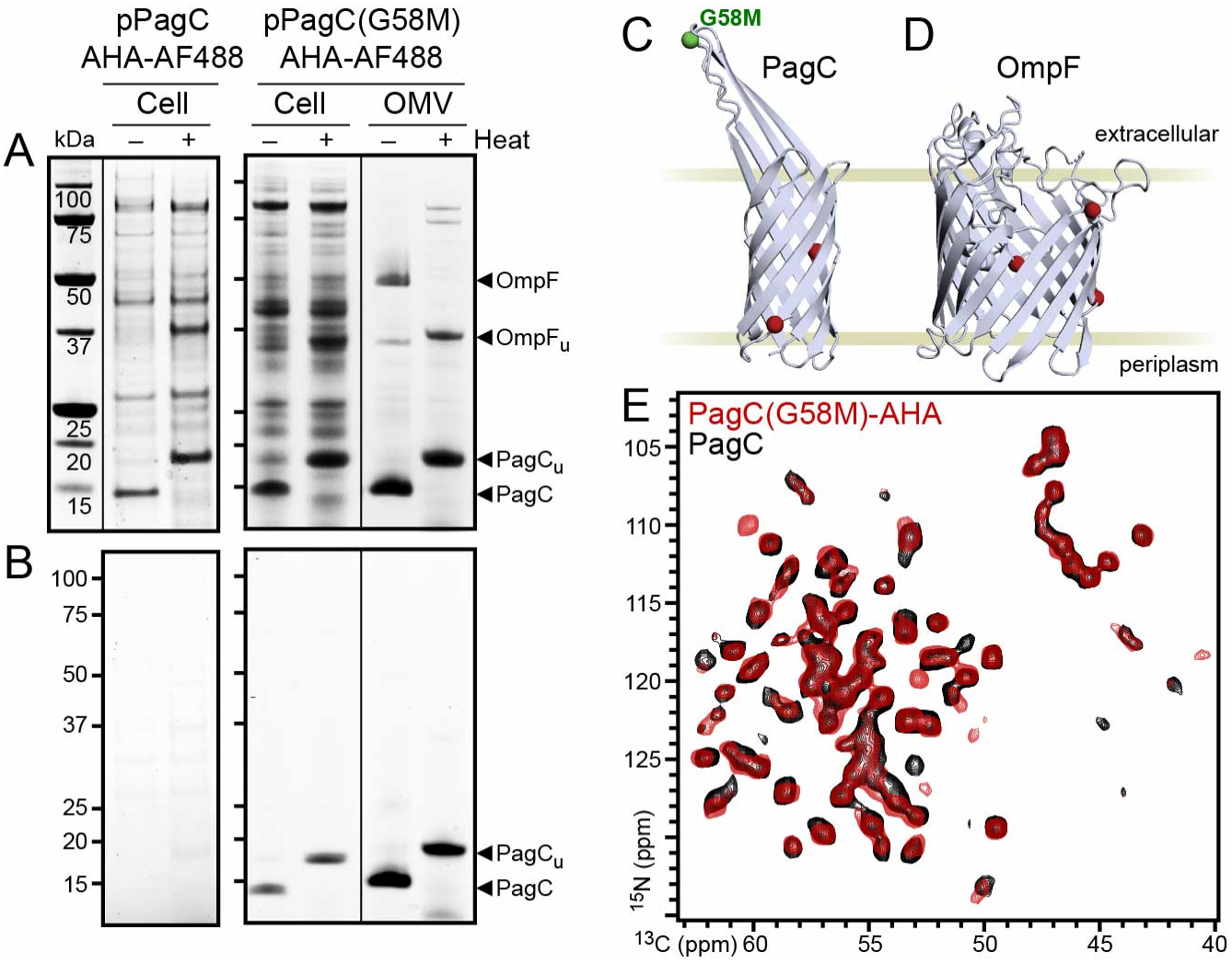
BONCAT labeling at the OM surface. **(A, B)** SDS PAGE of total cells or OMVs after CuAAC with fluorescent AF488-alkyne. The gels were developed with Coomassie stain (A) or imaged with AF488 fluorescence (B). **(C, D)** Structural models of PagC (from MD simulation ^6c^) and OmpF (from the crystal structure, PDB 2ZFG ^10^) showing CA atom positions (sphere) of native (red) or mutant G58M (green) Met residues. Only one monomer of the OmpF trimeric porin is shown. **(E)** Solid-state ^15^N/^13^C 2D NCA NMR spectra of AHA labeled wild-type pPagC (black) or mutant pPagC(G58M) (red) in OMVs.

BONCAT-mediated surface labeling of *E. coli* cells was first demonstrated for the major OM porin OmpC ^11^, and advances in reagent technology, including development of the hydrophilic THPTA triazole ligand ^12^, have improved reaction efficiency and specificity for the cell surface. Because AF488-alkyne is built on a hydrophilic, sulfonated rhodamine core carrying multiple ionic charges, it cannot passively diffuse across intact live-cell plasma membranes, and its molecular weight (572 Da) and physical size are very close to the 600-700 Da exclusion limit of the OmpF pore.

To explore PagC-specific labeling of at the OM surface, we introduced a G58M mutation at the apex of the second extracellular loop (**Fig. 2C, green**), then cultured the cells following protocol V, and subjected both intact cells and OMVs to CuAAC with AF488-alkyne.

SDS-PAGE shows that the G58M mutant protein is highly produced in both fractions, but while whole cells also contain many other proteins, and OMVs are primarily enriched in OmpF (**Fig. 2A**), the CuAAC reaction yields only pPagC-specific AF488 fluorescence with negligible background (**Fig. 2B**). Thus we conclude that while all proteins incorporate AHA at Met, pPagC(G58M)-AHA offers the only surface-accessible site for AF488-alkyne reaction.

Importantly, AHA tagging does not appear to perturb the overall β-barrel structure of the protein. On SDS PAGE, pPagC(G58M)-AHA retains the same heat modifiability property of wild-type pPagC: Folded at room temperature (~15 kDa) and unfolded with heat (~18 kDa) (**Fig. 2A, B**). Moreover, the 2D ^15^N/^13^C NCA solid-state NMR spectra of pPagC(G58M)-AHA and wild-type pPagC are nearly identical except for a few select ^15^N/^13^C chemical shift differences, pointing to local conformational and/or environmental changes in the vicinity of AHA-substituted G58M, M73 and M129. Overall, the data reflect only minimal system perturbation supporting the use BONCAT for functional OM protein studies.

To further visualize pPagC-specific labeling at the OM surface, we imaged the AF488-labeled pPagC(G58M) cells using structured illumination microscopy. In the absence of AHA supplementation, only minimal AF488 background is observed, likely arising from AF488-alkyne reactivity of occasional dead cells (**Fig. 3A, green**). By contrast, with AHA supplementation, the data reveal a shell of PagC(G58M)-AHA-AF488 fluorescence (**Fig. 3B, green**) co-localizing with the OM marker lipid stain (**Fig. 3, magenta**), and encapsulating the DNA marker Hoechst stain (**Fig. 3, blue**).

**Figure 3.**
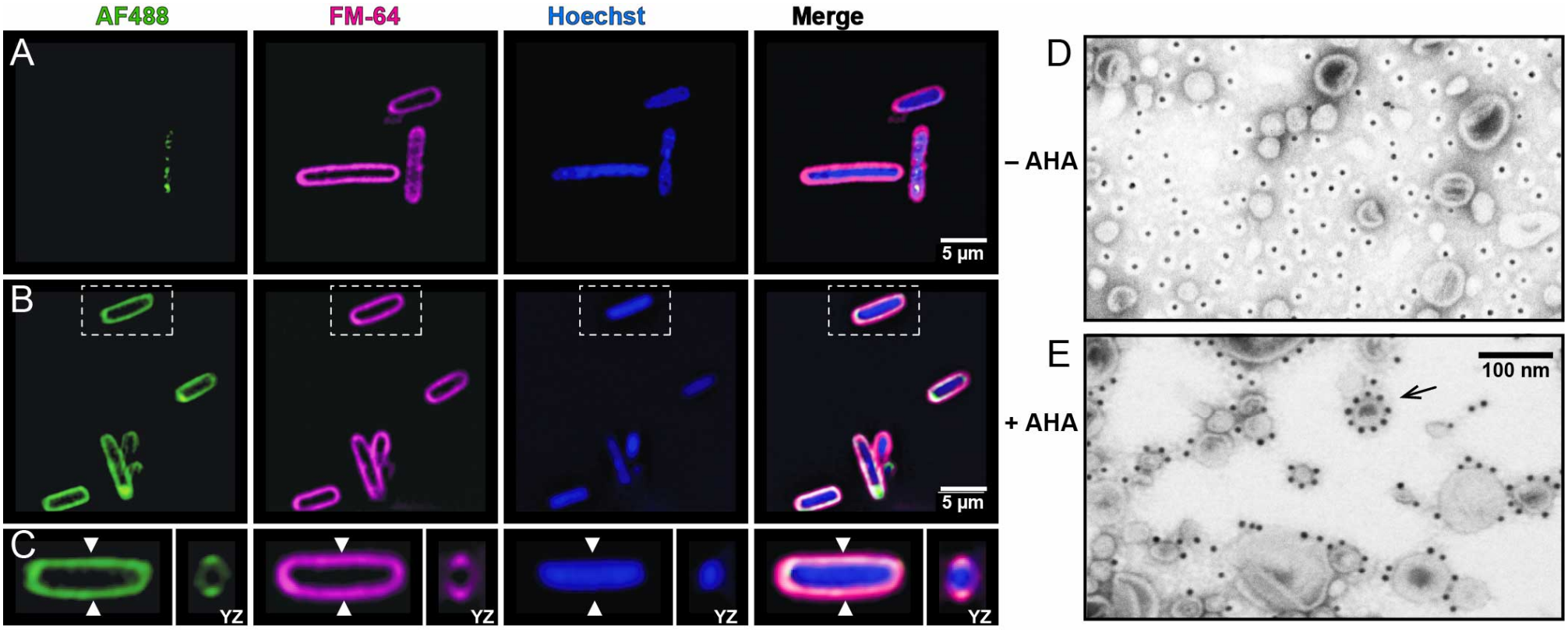
Surface labeling of PagC in whole cells and OMVs. Cells and OMVs were derived from cultures supplemented with Met (A, D) or AHA (B, C, E) during expression, and then subjected to CuAAC with AF488-alkyne (A-C) or 10 nm nano-Au-alkyne (D, E). **(A-C)** Fluorescence microscopy of pPagC(G58M) *E. coli* cells. Cells were AF488 labeled (green) and co-stained for lipid (pink; FM4-64) and DNA (blue; Hoechst) prior to imaging. Inset dashed line boxes (B) mark cells selected for z-stack analysis and white arrows designate YZ plane positions (C). **(D, E)** Negative stain electron microscopy of pPagC(G58M) OMVs. Black arrow points to a 30 nm OMV decorated with 9 nano-Au particles around its circumference.

Finally, we examined PagC-specific surface labeling with gold nanoparticles. Negative stain electron microscopy (EM) shows that alkyne-nano-Au (10 nm diameter; Sigma) associates specifically with the surface of pPagC(G58M) OMVs produced with AHA supplementation (**Fig. 3E**), but fails to associate with the OMV surface in the absence of AHA (**Fig. 3D**). In one example (**Fig. 3E, black arrow**), nine nano-Au particles appear to decorate the ~ 94 nm circumference of a ~30 nm diameter OMV, indicating dense and uniform distribution of PagC at the vesicle surface. Considering that the ~2.5 nm diameter of the PagC β-barrel may be expected to occupy a surface area of ~6.25 nm^2^, we estimate that the 10 nm nano-Au particles used in this study will only be able to access and label one PagC in an array of 4 or 5 neighboring molecules. The data suggest that this approach, combined with the use of smaller nano-Au particles, will shed light on OM protein localization and clustering in the bacterial OM. Taken together, the data demonstrate efficient surface CuAAC labeling of pPagC in both cells and OMVs and highlight the utility of this protocol for ultrastructural studies.

In conclusion, metabolic incorporation of the azide-functionalized Met-analog AHA, coupled with biorthogonal CuAAC with AF-488-alkyne, provides an effective platform for screening target-specific isotope labeling protocols for *in situ* NMR experiments of OM proteins. In this context, the BONCAT approach requires less extensive genetic modifications compared to either codon reassignment ^13^ or Amber codon suppression ^14^. The assay is rapid and expands the range of cell culture conditions that can be tested to achieve target specificity, bypassing the need for more intensive and costly NMR screening experiments.

The optimized labeling protocol developed with this rapid screening approach enables high-resolution NMR spectra to be obtained *in situ* with negligible off-target background signal from endogenous proteins. Moreover, when applied to whole cells and OMVs, biorthogonal CuAAC further enables target-specific, fluorescence or nano-Au tagging of proteins at the OM surface, extending the utility of the protocol for structure-activity studies of membrane proteins *in situ*.

## METHODS

### DNA, bacterial strains and culture conditions

All plasmids were cloned into the *E. coli* parent strain Lemo21(DE3) (New England BioLabs, MA, USA). The DNA fragments encoding PagC or PagC(G58M) were obtained by Genscript (Piscataway, NJ, USA) and designed for in-frame insertion into the pET22b(+) plasmid, downstream from the PelB signal peptide ^15^. Induction was confirmed by SDS-PAGE. Ampicillin and chloramphenicol were each utilized at 100 µg/mL and 35 µg/mL, when appropriate. Bacterial strains were cultured in M9 minimal media supplemented with 1 g/L (NH_4_)_2_SO_4_ and 0.2% glucose unless otherwise stated.

### Metabolic labeling with AHA

Cells from an overnight culture were sub-cultured into M9 media supplemented with 100 µM L-Met, and grown at 37°C up to a density corresponding to OD_600_~0.6. After cooling to room temperature (~20°C), cells were induced with IPTG for 10 min, then isolated by centrifugation, washed once with phosphate buffer saline (PBS), and resuspended in M9. For metabolic labeling (protocols II–V), AHA (Sigma Aldrich) was supplemented to 100 µM during the labeling window. During the starvation pulse (protocol V), cells were resuspended in M9 without glucose, ammonium or AHA, and these components sterile filtered into the medium after the starvation window.

### AF488 labeling of cell and OMV lysates

Cells were isolated by centrifugation, adjusted to a density of OD_600_=1 in a volume of 1 mL, washed with PBS, resuspended in 100 µL of BPER Complete (Invitrogen), frozen, and then mixed 1:1 with 2x lysis buffer (100 mM HEPES, 100 mM NaCl, 2% SDS, pH 7). OMVs were isolated by culture media ultrafiltration (100 kDa cutoff spin concentrator) followed by centrifugation as described ^5^, and then 200 µg of OMVs were mixed with 160 µL of 1x lysis buffer, heated to 95°C for 10 min, and centrifuged (12,000 x g, 2 min) to remove insoluble material.

For click reaction, 160 µL of each cell or OMV lysate sample was transferred to a clean tube, then supplemented with 2 µL of 2 mM AF488-alkyne (Invitrogen) and vortexed briefly to mix.

Separately, 10 µL of 100 mM Tris-(3-hydroxypropyltriazolylmethyl)-amine (THPTA, Sigma) were mixed with 2 µL of 100 mM Cu(II)SO_4_. The 12 µL mixture was incubated at room temperature for 5 min, added to the lysate sample and vortexed briefly to mix, and then supplemented with 10 µL of 100 mM sodium ascorbate and vortexed briefly to mix.

The reaction was gently rotated for 30 min in the dark, and then terminated by sequential addition of 600 µL methanol, 150 µL chloroform, and 400 µL ultrapure water with thorough vortexing in between. After removing the organic upper phase by centrifugation (16,000 x g, 5 min), the insoluble protein fraction was washed with 500 µL methanol, then again isolated by centrifugation and air dried before resuspending in 100 µL of fluorescence-compatible sample buffer (Invitrogen) for analysis.

### Surface labeling of live cells and intact OMVs

Cells were isolated by centrifugation, adjusted to a density of OD_600_=1 in a volume of 1 mL, then washed with PBS, and resuspended in 160 µLof 1x Dulbecco’s PBS (dPBS). OMVs were isolated and resuspended in dPBS. A total of 200 µg of OMVs were used for the reaction.

The pre-mixed THTPA and Cu(SO_4_) click reaction components were added, then supplemented with 10 µL of 100 mM sodium ascorbate, and the reactions allowed to proceed in the dark for 30 min as described above. For AF488 surface labeling, the click reaction was performed with 2 µL of 2 mM AF488-alkyne. For gold labeling, 8 µL of 10 nm, alkyne-functionalized gold nanoparticles (Sigma 928070) were used in the reaction.

For SDS-PAGE analysis, cells were isolated by centrifugation, washed three times with dPBS, and resuspended in 400 µL of PBS, and then mixed 1:1 with 4x fluorescence-compatible sample buffer. OMVs were washed three times and resuspended in OMV buffer (50 mM HEPES pH 7, 0.9 mM CaCl_2_, 0.5 mM MgCl_2_) by ultrafiltration with a 100 kDa cutoff spin concentrator, and 3 µg of labeled OMVs were mixed with SDS sample buffer.

### Fluorescence detection of SDS-PAGE

Samples were separated on a 4-12% NuPAGE gradient gel (Invitrogen) and imaged with a Bio-Rad Chemidoc using the SYBR green channel with autoexposure. Gels were stained with Coomassie to detect total protein. The AF488 fluorescence intensity in each lane was measured in FIJI with the gel analyzer plugin, and the intensities then used to calculate pPagC signal intensity as percent of total signal.

### Fluorescence microscopy

Following click reaction, the washed cells in dPBS were supplemented with 1 µg/mL of FM4-64 dye (Invitrogen) and 1 µg/mL of Hoechst 33324 dye (Invitrogen), and then adjusted to a density of OD_600_ ~ 0.2. For microscopy, 5 µL of cell suspension were spread onto a freshly prepared 9 mm pad of 1% low-melt agarose, supplemented with 1x M9 salts, 1 µg/mL FM4-64 dye, and 1 µg/mL Hoechst 33324, and dye cast in a press-to-seal silicone isolator (Invitrogen) on a glass slide.

The sample was dried on the agarose pad for 5 min, then covered with a #1.5 glass coverslip, and sealed with clear nail polish. Samples were imaged immediately on a Nikon N-SIM microscope equipped with a 100x oil objective and an MCL NanoDrive PiezoZ Drive. For each image, a Z stack was collected using 0.2 µm steps and reconstructed using the Nikon software.

Exposure settings were determined on the AHA-positive samples and used for the AHA-negative controls for direct comparison. Maximum intensity projections and orthogonal views were generated using FIJI software ^16^.

### Electron microscopy

Gold-labeled OMVs were adsorbed onto glow-discharged 400 mesh Formvar carbon copper grids (Electron Microscopy Sciences) for 15 sec, and the grids then stained with 2% uranyl-acetate for 30 sec before imaging on a Jeol JEM-1400 Transmission Electron Microscope (JEOL USA, MA) at 80 kV.

### Solid-state NMR

OMVs were centrifuged directly into 3.2 mm zirconium MAS rotors, and NMR experiments were performed on a 700 MHz AVANCE NEO Bruker spectrometer equipped with a 3.2 mm BlackFox probe. Spectra were acquired at a sample temperature of 4°C, with a MAS rate of 12.5 kHz, and a recycle delay of 2 s. The 1D ^15^N spectra were acquired with a ^1^H-^15^N CP contact time of 1 ms and 10,000 (pEV) or 512 (pPagC) transients. The 1D ^13^C spectra were acquired with CP contact times of 1 ms (^1^H-^13^C CP) and 4 ms (^15^N-^13^C) and 512 transients.

The 2D ^15^N/^13^C NCA spectra were recorded using Hadamard encoding ^17^ with 512 transients and 100 t1 increments, for a total experimental time of 28 hours. Other experimental settings were as described ^5^.

## ACKNOWLEDGEMENTS

This work was supported by grants from the National Institutes of Health (NIH) including research grants (GM118186 and F32AI188770) and a shared instrumentation grant for NMR instrumentation (OD028716). All imaging was completed at the Oxford Instruments Center for Advanced Microscopy – Electron Microscopy Core at the Medical College of Wisconsin (RRID:SCR_026315). We thank Ashish Gadicherla and Meghan Hardwick for their assistance with microscopy, Akanksh Gopinath for assistance with assay development, and Galia Debelouchina and Kyungsoo Shin for insightful discussion.

## ABBREVIATIONS

AF488: AlexaFluor-488
BONCAT: bioorthogonal non-canonical amino acid tagging
CuAAC: copper-catalyzed azide-alkyne cycloaddition
OM: outer membrane
OMV: outer membrane vesicle

## AUTHOR CONTRIBUTIONS

NAW, TG and FMM were involved in research design, data analysis and discussion. NAW and TG performed experiments. NAW and FMM wrote the manuscript with input and approval from all authors.

## REFERENCES

(1) Marassi, F. M.; Pintacuda, G. Curr. Opin. Struct. Biol. 2025, 94, 103129.

(2) (a)Hartmann, G.; Honikel, K. O.; Knusel, F.; Nuesch, J. Biochim Biophys Acta 1967, 145, 843–844. (b)Tabor, S.; Richardson, C. C. Proc. Natl. Acad. Sci. U. S. A. 1985, 82, 1074–1078.

(3) (a)Lee, K. M.; Androphy, E. J.; Baleja, J. D. J. Biomol. NMR 1995, 5, 93–96. (b)Almeida, F. C.; Amorim, G. C.; Moreau, V. H.; Sousa, V. O.; Creazola, A. T.; Americo, T. A.; Pais, A. P.; Leite, A.; Netto, L. E.; Giordano, R. J.; et al. J Magn Reson 2001, 148, 142–146. (c)Cruzeiro-Silva, C.; Albernaz, F. P.; Valente, A. P.; Almeida, F. C. Cell Biochem. Biophys. 2006, 44, 497–502. (d)Narasimhan, S.; Pinto, C.; Lucini Paioni, A.; van der Zwan, J.; Folkers, G. E.; Baldus, M. Nat Protoc 2021, 16, 893–918.

(4) (a)Renault, M.; Tommassen-van Boxtel, R.; Bos, M. P.; Post, J. A.; Tommassen, J.; Baldus, M. Proc. Natl. Acad. Sci. U. S. A. 2012, 109, 4863–4868. (b)Castegnaro, F.; Burmann, B. M.; Thoma, J. Methods Mol. Biol. 2023, 2652, 247–257.

(5) Wood, N. A.; Kraft, A.; Shin, K.; Gopinath, T.; Marassi, F. M. Proc. Natl. Acad. Sci. U. S. A. 2025, 122, e2501638122.

(6) (a)Kitagawa, R.; Takaya, A.; Ohya, M.; Mizunoe, Y.; Takade, A.; Yoshida, S.; Isogai, E.; Yamamoto, T. J. Bacteriol. 2010, 192, 5645–5656. (b)Dehinwal, R.; Cooley, D.; Rakov, A. V.; Alugupalli, A. S.; Harmon, J.; Cunrath, O.; Vallabhajosyula, P.; Bumann, D.; Schifferli, D. M. mBio 2021, 12, e0086921. (c)Dehinwal, R.; Gopinath, T.; Smith, R. D.; Ernst, R. K.; Schifferli, D. M.; Waldor, M. K.; Marassi, F. M. Nat. Commun. 2024, 15, 6958.

(7) (a)Bernstein, J. A.; Khodursky, A. B.; Lin, P. H.; Lin-Chao, S.; Cohen, S. N. Proc. Natl. Acad. Sci. U. S. A. 2002, 99, 9697–9702. (b)Selinger, D. W.; Saxena, R. M.; Cheung, K. J.; Church, G. M.; Rosenow, C. Genome Res. 2003, 13, 216–223. (c)Chen, H.; Shiroguchi, K.; Ge, H.; Xie, X. S. Mol Syst Biol 2015, 11, 808. (d)Deighan, P.; Free, A.; Dorman, C. J. Mol. Microbiol. 2000, 38, 126–139.

(8) (a)Dieterich, D. C.; Link, A. J.; Graumann, J.; Tirrell, D. A.; Schuman, E. M. Proc. Natl. Acad. Sci. U. S. A. 2006, 103, 9482–9487. (b)Kiick, K. L.; Saxon, E.; Tirrell, D. A.; Bertozzi, C. R. Proc Natl Acad Sci U S A 2002, 99, 19–24.

(9) Hatzenpichler, R.; Scheller, S.; Tavormina, P. L.; Babin, B. M.; Tirrell, D. A.; Orphan, V. J. Environ Microbiol 2014, 16, 2568–2590.

(10) Yamashita, E.; Zhalnina, M. V.; Zakharov, S. D.; Sharma, O.; Cramer, W. A. The EMBO Journal 2008, 27, 2171–2180.

(11) (a)Link, A. J.; Tirrell, D. A. J. Am. Chem. Soc. 2003, 125, 11164–11165. (b)Link, A. J.; Vink, M. K.; Tirrell, D. A. J. Am. Chem. Soc. 2004, 126, 10598–10602.

(12) Chan, T. R.; Hilgraf, R.; Sharpless, K. B.; Fokin, V. V. Organic Letters 2004, 6, 2853–2855.

(13) (a)Kiick, K. L.; van Hest, J. C. M.; Tirrell, D. A. Angewandte Chemie International Edition 2000, 39, 2148–2152. (b)Link, A. J.; Mock, M. L.; Tirrell, D. A. Curr Opin Biotechnol 2003, 14, 603–609.

(14) (a)Noren, C. J.; Anthony-Cahill, S. J.; Griffith, M. C.; Schultz, P. G. Science 1989, 244, 182–188. (b)Bain, J. D.; Diala, E. S.; Glabe, C. G.; Wacker, D. A.; Lyttle, M. H.; Dix, T. A.; Chamberlin, A. R. Biochemistry 1991, 30, 5411–5421.

(15) Lei, S. P.; Lin, H. C.; Wang, S. S.; Callaway, J.; Wilcox, G. J. Bacteriol. 1987, 169, 4379–4383.

(16) Schindelin, J.; Arganda-Carreras, I.; Frise, E.; Kaynig, V.; Longair, M.; Pietzsch, T.; Preibisch, S.; Rueden, C.; Saalfeld, S.; Schmid, B.; et al. Nature methods 2012, 9, 676–682.

(17) Gopinath, T.; Kraft, A.; Shin, K.; Wood, N. A.; Marassi, F. M. J. Biomol. NMR 2025, 79, 35–45.

