## Supporting Information for "Rapid screening of target-specific isotope labeling protocols for *in situ* NMR of bacterial membrane proteins"

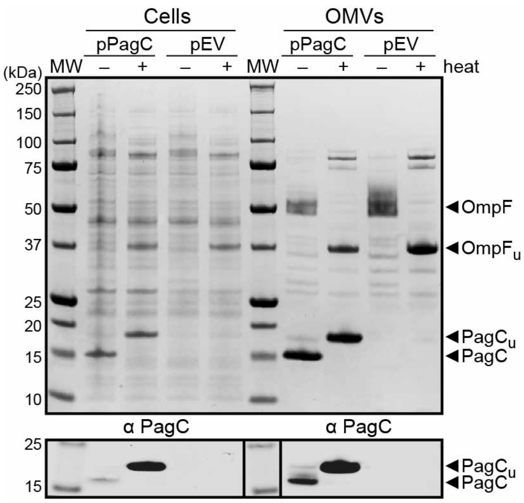


**Figure S1. SDS-PAGE of whole cells and OMVs isolated from pPagC or pEV bacteria (adapted from [S1]).** Proteins were visualized with Coomassie stain (top) or immunoblotting with PagC-specific antibody (α-PacC, bottom). Arrows mark bands from folded or heat-unfolded (u) proteins.

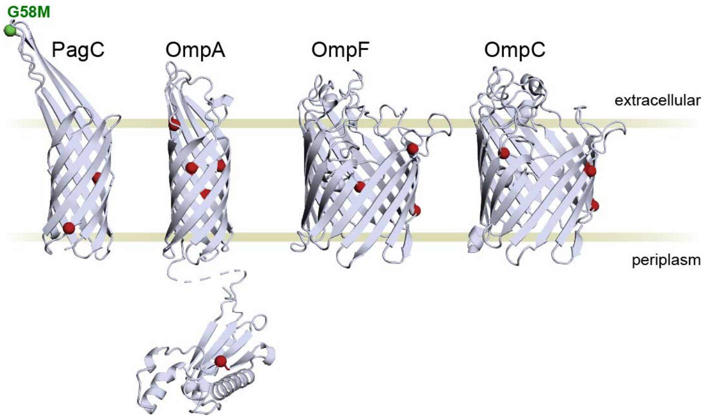


**Figure S2. Native Met residues in PagC and the major OM proteins.** CA atom positions (sphere) denote native Met residues (red) or mutant G58M (green). Structural models represent PagC (from MD simulation [S2]), OmpA (AlphaFold, AF-P0A910), OmpF (crystallography, PDB 2ZFG [S3]) and OmpC (crystallography, PDB 2J1N [S4]). Only one monomer is shown for each OmpF and OmpC trimeric porin for simplicity.

S1. Wood NA, Kraft A, Shin K, Gopinath T, Marassi FM (2025) In situ NMR reveals a pH sensor motif in an outer membrane protein that drives bacterial vesicle production. Proc. Natl. Acad. Sci. U. S. A., 122, e2501638122. (PMC12232643)

S2. Dehinwal R, Gopinath T, Smith RD, Ernst RK, Schifferli DM, Waldor MK, Marassi FM (2024) A pH-sensitive motif in an outer membrane protein activates bacterial membrane vesicle production. Nat. Commun., 15, 6958. (PMC11322160)

S3. Yamashita E, Zhalnina MV, Zakharov SD, Sharma O, Cramer WA (2008) Crystal structures of the OmpF porin: function in a colicin translocon. The EMBO Journal, 27, 2171–2180.

S4. Baslé A, Rummel G, Storici P, Rosenbusch JP, Schirmer T (2006) Crystal Structure of Osmoporin OmpC from E. coli at 2.0 Å. J. Mol. Biol., 362, 933–942.

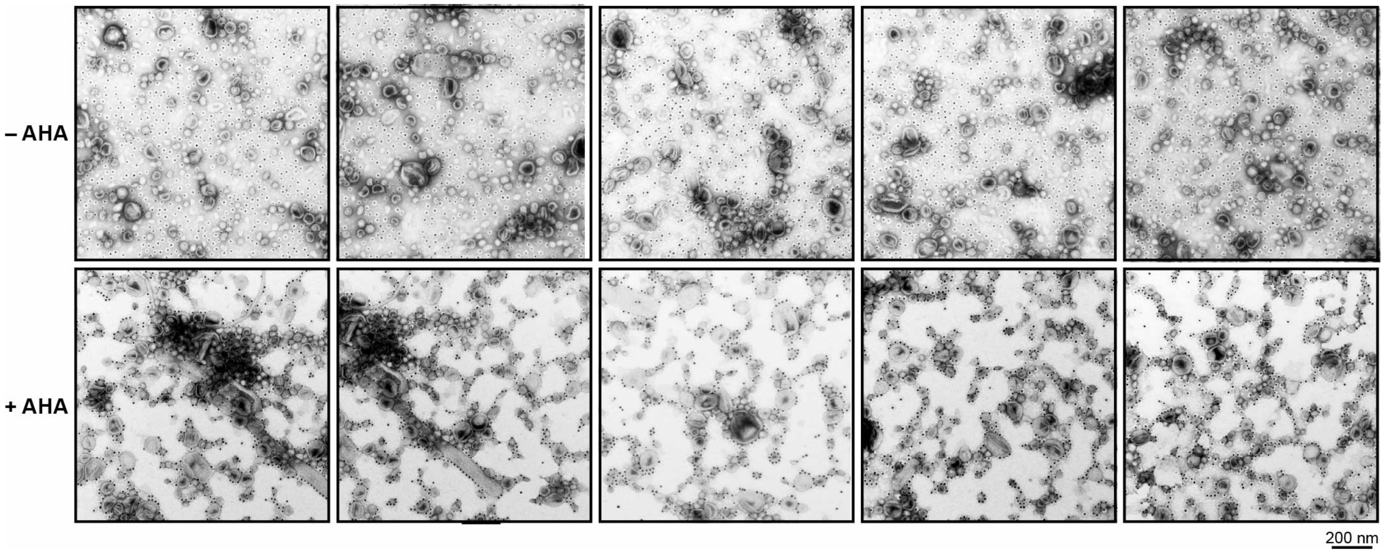


**Figure S3. Targeted nano-Au labeling at the OMV surface.** Negative stain electron microscopy of PagC(G58M) OMVs from cultures supplemented with Met (–AHA) or AHA (+AHA) and then subjected to CuAAC with 10 nm nano-Au-alkyne.
